# Generation of transgenic quails by *in vivo* transfection of primordial germ cells

**DOI:** 10.1101/625665

**Authors:** Olivier Serralbo, Nadège Véron, Caitlin Cooper, Marie-Julie Dejardin, Timothy Doran, Christophe Marcelle

**Affiliations:** EMBL Australia; Australian Regenerative Medicine Institute (ARMI), Monash University, Building 75, Clayton, Victoria 3800, Australia; Institut NeuroMyoGène (INMG), University Claude Bernard Lyon1, CNRS UMR 5310, INSERM U1217, Lyon, France; CSIRO Health & Biosecurity, Australian Animal Health Laboratory, Geelong Vic, Australia

## Abstract

During development, dynamic changes in tissue shapes known as morphogenesis result from the exquisite orchestration of signalling pathways, cell–cell interactions, cell divisions, and coordinated movements. How cells within embryos adopt a vast array of cell fates and tissue shapes in such an ever-changing environment has fascinated scientists for generations, yet the ability to observe and characterize those rapid changes has proven technically challenging in higher vertebrates. The japanese quail (*Coturnix coturnix japonica*) is an attractive model where basic rules driving morphogenesis in amniotes can be deciphered using genetic approaches. Similar to the more popular chicken model, the quail embryo is easily accessible to a wide range of manipulations and live imaging. A decisive asset of quail over chicken is a much shorter life cycle, which makes its use as genetic model for basic research extremely appealing. To date, all existing transgenic quail lines were generated using replication-deficient lentiviruses, but diverse limitations of this approach have hindered the widespread expansion of such technology. Here, we successfully used a plasmid-based *in vivo* transfection of quail primordial germ cells (PGCs) to generate a number of transgenic quail lines over a short period of time. The plasmid-based approach is simple, efficient and it allows using the infinite variety of genome engineering approaches developed in other models, such as strategies to facilitate transgenic bird selection, shown here. This major technological development completes the vast panel of techniques applicable to the avian model, making it one of the most versatile experimental systems available.

## Introduction

Due to the easy access of chicken embryos to manipulation and high-end imaging, this model has been at the origin of numerous seminal discoveries in a diverse range of topics (e.g. immunology, genetics, virology, cancer, cell biology; Stern, 2005). However the production of transgenic chicken has lagged behind the two main genetic vertebrate models (mouse and fish), largely due to the specificities of the reproductive physiology of birds. The zygote is very difficult to access as it develops internally in the hen’s oviduct and on a large yolk. By the time the egg is laid, the embryo has already developed into a blastoderm of about 40,000-50,000 cells (Eyal-Giladi and Kochav, 1976; Intarapat and Stern, 2013). Because of this, most researchers in avian genetics have focused their efforts on two distinct methods (Nishijima and Iijima, 2013): i) the genetic manipulation of primordial germ cells (PGCs) *in vitro*, which are injected back into recipient embryos (Park et al., 2014) or ii) the infection of the blastoderm with replication-defective lentiviruses (Bosselman et al., 1989; McGrew et al., 2004). Both approaches have been applied successfully to chicken (Nishijima and Iijima, 2013). However, due to the long-life cycle of chicken (6 months to sexual maturity) and the relatively low efficiency and high technicality of those technologies, they have not been widely used in basic research and their development has been mainly driven by industrial interests to produce biologically active pharmaceutical proteins in eggs (Lillico et al., 2007; Nishijima and Iijima, 2013; Woodfint et al., 2018).

Japanese quail (*Corturnix coturnix japonica*) is a better alternative for avian genetics. Quail is a well known model in the developmental biology field through its extensive use in the so-called quail-chick chimera technique (Le Douarin, 1973, 2005). Indeed, quail and chicken are very close relatives (Order: Galliformes; Family: Phasianidae), they share on average 95% homology at the gene sequence level (GenBank; see accompanying paper) and quails are susceptible to most of the chicken diseases (Barnes, 1987). Similar to chicken, quail lay about one egg per day, however their smaller size makes them more compatible with the frequently limited space of animal facilities. A decisive advantage of quail over chicken is that they reach sexual maturity in 6 weeks. Thus, their life cycle is significantly shorter than that of chicken (26 weeks), but also of mice (8 weeks) or zebrafish (12 weeks). As no reliable technique for the culture of quail PGCs has yet been described, transgenesis in this species has mainly relied on the infection of blastoderm cells (or circulating PGCs, Zhang et al., 2012) with lentiviruses carrying fluorescent markers under ubiquitous or tissue-specific promoters, leading in recent years to the generation of 7 quail lines: ubiquitous (Huss et al., 2015; Zhang et al., 2012); neuronal (Scott and Lois, 2005; Seidl et al., 2013); adipose (Ahn et al., 2015); intestinal (Woodfint et al., 2017); endothelial (Sato et al., 2010). A high titre of lentivirus is required to succeed at producing chimeric embryos carrying transgenic PGCs using this technique (Poynter and Lansford, 2008; Scott and Lois, 2005). Even though large inserts (more than 10 kilobases) can be packaged using lentiviruses, a sharp drop in viral titre is observed with insert above 4-5 kb, likely representing the upper limit of constructs that can be used to generate transgenic birds (Kumar et al., 2001; our observation). This may be sufficient for a number of applications, but the size limitation is a significant restriction in the use of lentiviruses to generate complex transgenes.

For biosafety and commercial reasons, non-viral and simpler technologies for achieving transgenic poultry have been tested. The direct *in vivo* transfection of PGCs with commercially available liposome-based transfection reagents has recently been successfully used to generate transgenic chicken lines (Cooper et al., 2018b, 2018a; Tyack et al., 2013). Insertion of Tol2 transposable elements in constructs allows the efficient and stable integration of foreign DNA into the genome of transfected PGCs in the presence of exogenously provided transposase protein. Since DNA insert up to a size of 11kb can be cloned between Tol2 sequences with no visible loss of transposable efficiency (Kawakami, 2007) this allows using larger and/or more complex constructs than in lentiviruses. Another advantage of transgenesis using the Tol2 transposon system is that it leads to minimal epigenetic silencing during development (Macdonald et al., 2012).

Here we used the direct transfection of PGCs in the bloodstream of quail embryos. We present three novel quail transgenic lines carrying fluorescent proteins under ubiquitous and tissue-specific promoters. Illustrating the flexibility of the plasmid-based system, we integrated a cassette containing the GFP protein driven by a promoter for Chick δ1-Crystallin (Reneker et al., 2004) making the identification of transgenic animals possible at hatching by UV illumination (with UV goggles).

## Results

### Direct transfection of quail PGCs

Unlike mammals, avian PGCs temporarily transit through the blood system. Chicken PGCs initially located at the end of gastrulation within the germinal crescent, enter the extra-embryonic blood vessels at about 30 hours of incubation (E1.5, HH9) and begin to circulate throughout the embryo. Their number in the blood peaks around E2/2.5 (HH15-16). By E3 (HH20), PGCs actively migrate back in the embryo and into the gonad anlagen (Nieuwkoop and Sutasurya, 1979). Chicken PGCs are transfected during their transient journey in the bloodstream, using a transfection mix contains lipofectamine, a transgenesis plasmid with Tol2 elements flanking the DNA to be inserted and a plasmid coding for transposase under an ubiquitous promoter (Tyack et al., 2013).

To test whether the direct transfection of PGCs can be achieved in quails, we determined whether we could observe PGCs transfected with fluorescent proteins once they colonized the gonads. A transfection mix containing a transgenesis plasmid coding for a membrane-localised GFP (GFPcaax) and a nuclear monomeric Cherry (nls-mCherry) under the control of a strong ubiquitous promoter (CAGGS: CMV promoter, chick beta actin enhancer; Niwa et al., 1991) was injected into the bloodstream of quail embryos (Fig1 A). Quail develop slightly faster than chicken (17-18 days for quails; 21 days for chicken). However, we found that the timing of injection of the transfection mix resulting in the colonization of fluorescent PGCs was similar to that determined in chicken, i.e. at 2 days (E2) of quail embryonic development. Indeed, 5 days after injection (at E7), we observed that gonads contained many fluorescent cells (Fig1 B). We confirmed that these were PGCs using whole-mount immunostaining with a PGC-specific marker VASA (Fig1 C-F).

**Figure 1.**
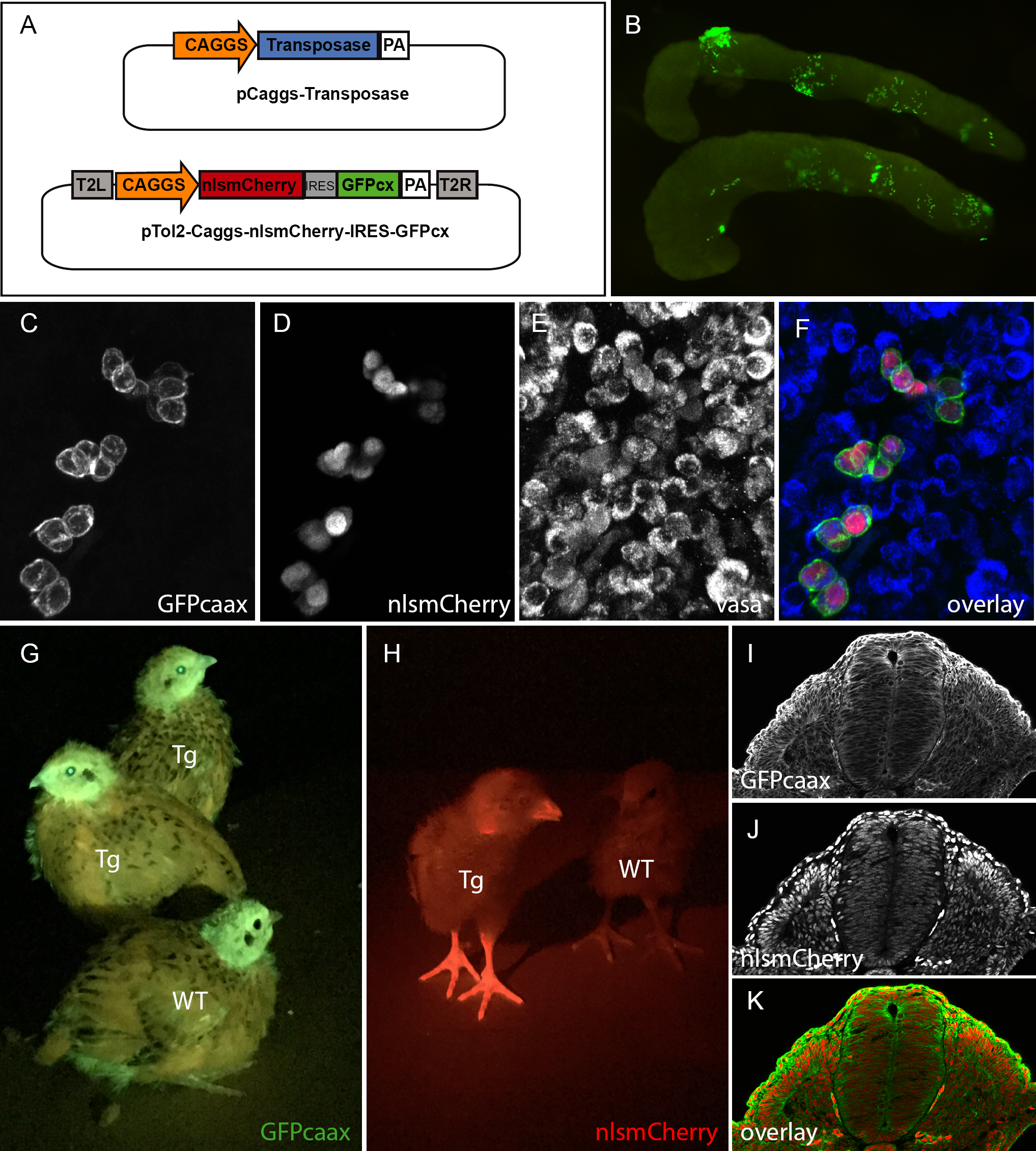
*In vivo* PGC transfection by direct injection. A) vectors used in the injection mix. B) Gonads from E7 embryo dissected 5 days after injection showing GFP positive PGCs. (C-F) Confocal view of transfected PGCs form E7 injected embryo expressing the PGC marker Vasa. (C) GFP, (D) nlsmCherry, (E) Vasa marker of PGCs (F) merge. (G-H) 14 day old transgenic quail (Tg) and wild type (WT) showing ubiquitous expression of GFP and mCherry when exposed to the appropriate wavelength light using UV goggles. (I-J) cross-section of an E3 transgenic embryo showing the ubiquitous expression of the transgene and the cellular localization of GFPcaax and nlsmCherry.

### Generation of a lens-specific GFP minigene to facilitate the selection of transgenic birds

To facilitate the selection of transgenic (F1) birds, we devised a fluorescent selection marker readily visible in the lens at hatching under UV illumination. We isolated a 462bp-long lens-specific promoter of the βB1crystallin gene (Duncan et al., 1995) from chicken genomic DNA and cloned it upstream of GFP to develop a selection mini-gene (CrystallGFP) based on lens expression (Fig2 H). To test the specificity of the promoter, we co-electroporated the CrystallGFP construct together with an ubiquitously expressed RFP (CAGGS-RFP) into the optic cup of an E3 embryo (Fig2 A-D). Twenty-four hours after electroporation, RFP-positive cells were found in the retina and lens (Fig2 B). However, only lens cells expressed the GFP (Fig2 C-D), showing the specificity of the βB1crystallin promoter for lens tissues. Transgenic embryos carrying the CrystallGFP minigene displayed a strong and specific expression of GFP in the lens (Fig2 E-G), while adult transgenic quail can be easily selected using UV goggles for GFP (Fig2 I). The CrystallGFP selection cassette was included in some of the constructs described below.

**Figure 2.**
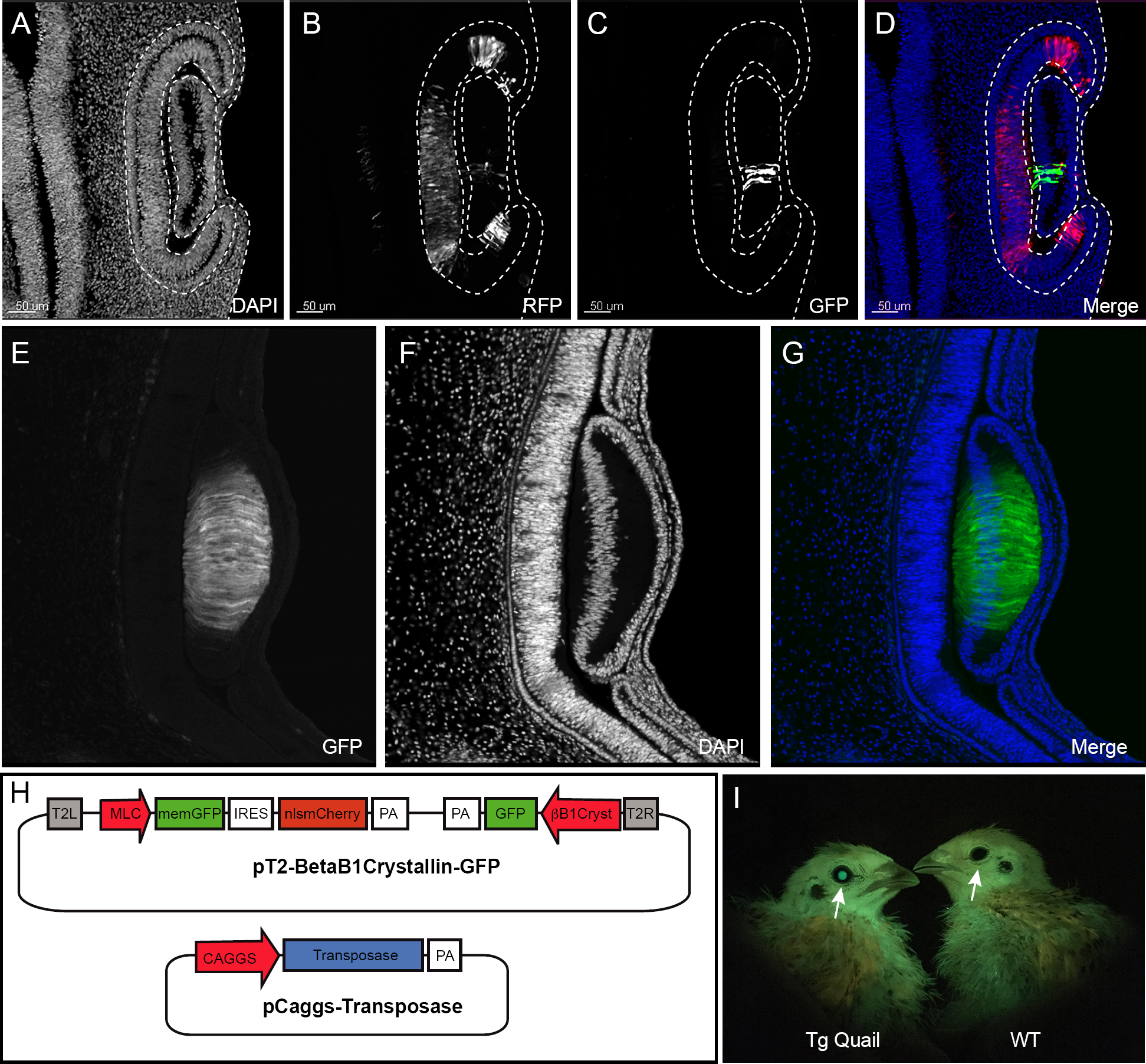
Design of CrystallGFP minigene. (A-D) cross-section an electroporated E3.5 embryo to test the specificity of the Crystallin promoter. (A) Dapi, (B) electroporation marker Caggs-RFP plasmid, (C) GFP, (D) Overlay. (E-G) Cross-section of an E5 transgenic embryo showing the specific expression of GFP in all lens cells (E) GFP, (F) DAPI, (G) Merge. (H) muscle-specific expression construct containing the betaB1 crystalin-GFP mini gene and co-injected with the transposase plasmid. (I) Muscle-specific 10 day old transgenic quail showing the specific GFP expression in lens (arrow) compared to wild type allowing an easy selection of transgenic birds by using GFP goggles.

### Generation of transgenic quails lines

#### CAGGS GFP-caax/nls-mCherry: ubiquitous expression of membranal GFP and nuclear RFP

To generate a transgenic quail line ubiquitously expressing a membrane-bound GFP and a nuclear RFP, embryos injected with the CAGGS GFP-caax/nls-mCherry plasmid (see above) were incubated until hatching and raised to adult stage. In this and other experiments described below, we have observed that about half of the (50) injected eggs hatched. Six weeks later, we collected semen from adult males and tested by PCR for the presence of the transgene. Three (F0) males that showed a positive PCR band for the transgene were crossed with four females each. From these crosses, transgenic (F1) birds could be readily spotted at hatching as they ubiquitously expressed GFP and mCherry (Fig1 G,H). Expression of GFP or RFP was visible in the beak, eyes and legs of the transgenic birds compared to wild type animals (Fig1 G,H). Immunostaining on cross-sections of E3 transgenic embryos showed an ubiquitous expression of the GFP at the cell membrane and of mCherry in nuclei (Fig1 I-K). From this and other crosses we have performed in the laboratory (see below), we estimate that about 1% of the offspring contain the transgene, an efficiency comparable to that observed in the chicken using the same technology (Tyack et al., 2013). Compared to the existing quail lines carrying ubiquitously expressed fluorescent proteins, this line should prove useful to researchers in the field, since the membrane-bound GFP results in a better resolution of cell membrane processes (protrusions, filopodia, etc.) than a cytoplasmic counterpart, while it also combines a nuclear mCherry, allowing accurate segmentation of cells necessary for automated image analyses such as for 3D cell tracking.

### Skeletal muscle-specific reporter quail

We generated a line carrying a promoter for the mouse alkali Myosin Light Chain gene (MLC; Kelly et al., 1995) upstream of the membrane-bound GFP and the nuclear mCherry reporters described above. We designed a muscle-specific promoter, based on a synthetic reporter derived from the MLC1F/3F gene regulatory sequences previously utilized for mouse transgenesis (3F-nlacZ-E; Kelly et al., 1995). It contains a 2 kb sequence located 5’ and 3’ of the MLC3F transcriptional start site together with a 260 bp enhancer sequences from the 3’ UTR region of the MLC3F gene, necessary for the high level of transcription in muscles. This construct was shown to drive strong LacZ expression in all (head and body) striated muscles from the early steps of myogenesis in somites of mouse embryos throughout embryogenesis, as well as in all skeletal muscles of the foetus and in the adult (Kelly et al., 1995). Because muscle expression is not readily visible in adult animal, we included the CrystallGFP cassette in the construct to facilitate the selection of F1 transgenic birds.

F0 founder males were crossed with females and from 242 chicks that hatched, 3 transgenic F1 founders (1 male and 2 females, i.e. 1.2% efficiency) were selected, based on the expression of GFP in the lens (Fig2 F). These F1 were used to characterize the expression of the transgene during embryogenesis.

Transgenic embryos were selected using the CrystallGFP marker (Fig3 I-K, arrow). The GFP and RFP reporters were expressed in all skeletal muscles of the developing embryos. Native GFP and mCherry expressions were brightly visible under UV examination at E3 in trunk somites (arrowheads in Fig3 I-K), while all trunk, head and limb muscles were easily detected at E5 (Fig3 L-N). Expression of the two fluorescent proteins in the myotome of trunk somites was confirmed on sections of E3 embryos (Fig3 A-D). Interestingly, we observed that GFP expression was detected within the transition zone (TZ, Fig3 E-H), where cells emanating from the medial border of the overlying dermomyotome first translocate before extending towards the anterior and posterior borders of each somite (Gros et al., 2004). When cells enter the TZ, they have already initiated the myogenic program and express Myf5. Once in the TZ, these progenitors initiate Myogenin (MyoG) expression and only after they are fully extended into myocytes, they initiate Myosin Heavy Chain expression (MyHC; Rios et al., 2011; Sieiro et al., 2016; our observation). We conclude that the MLC promoter becomes active after Myf5, concomitant to MyoG and before MyHC. Our observations are entirely coherent with what was observed in mouse with a similar promoter construct. However, the expression of LacZ that was observed in non skeletal tissues (brain, optic vesicle and heart) in 3F-nlacZ-E mouse embryos (Kelly et al., 1995) was not detected in the transgenic quails we analyzed, suggesting a more rigorous restriction to the skeletal muscle lineage. Whole mount staining of E3, E5 and E7 embryos showed a strong and specific expression of all skeletal muscles of the body and the head.

**Figure 3.**
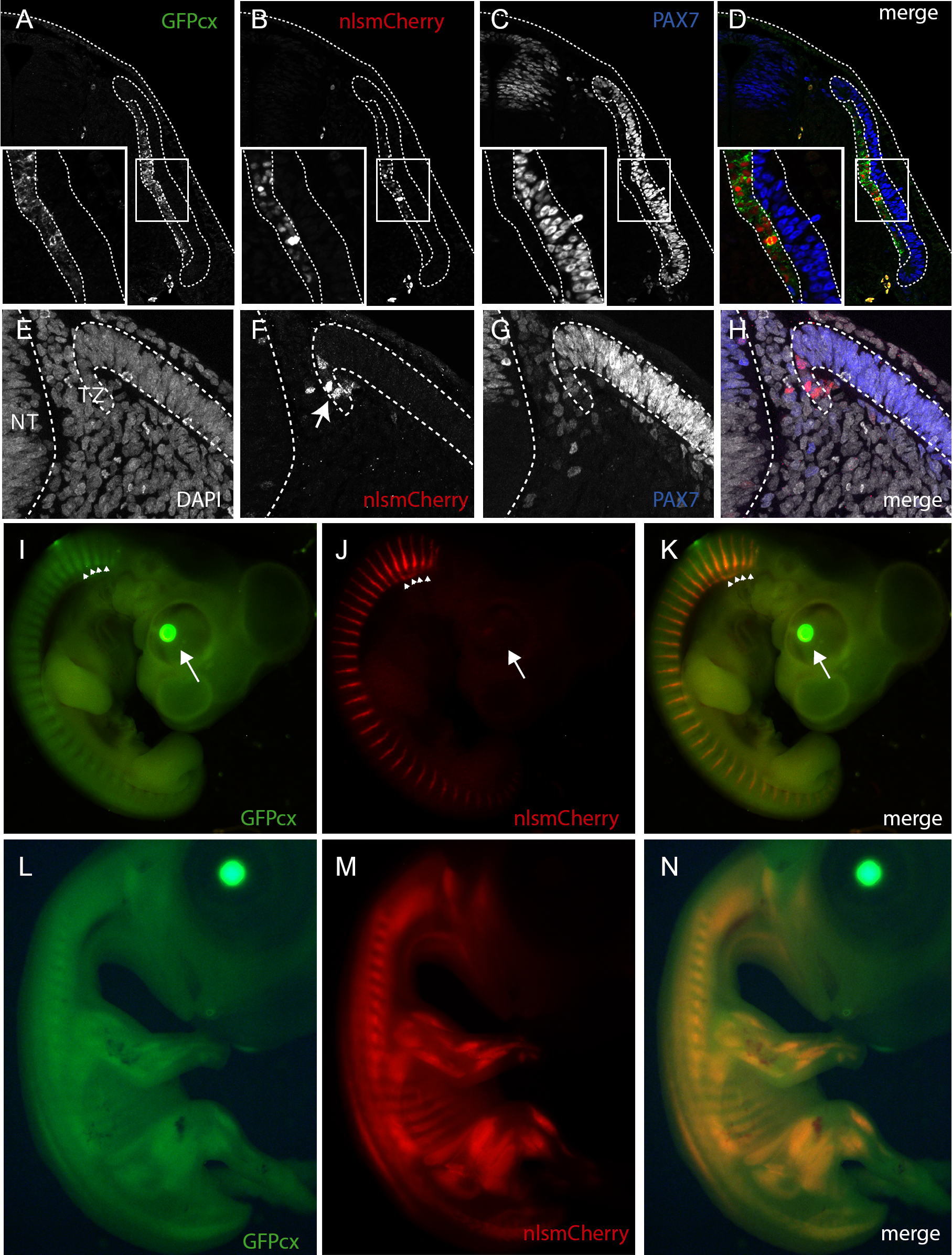
Description of muscle specific transgenic quail Tg(MLC-nlsmCherry-IRES-GFPcaxx). (A-D) cross section and immunostaining of an E3.5 transgenic embryo showing the expression of the transgene in the primary myotome. (A) GFPcaxx, (B) nlsmCherry, (C) Pax7, (D) overlay. A-D insert, magnification showing the membrane and nuclear localisation of GFP and mCherry, respectively, in primary myotomal cells. (E-H) Expression of the MLC transgene in the transition zone (TZ, F arrow), (E) DAPI, (F) mCherry, (G) Pax7. (I-K) E5 transgenic embryo showing native GFPcaax (E) and nuclear mCherry (F) expression in each myotome (arrowhead). Transgenic embryos can be selected with BetaB1 Crystallin expression (I, arrow). (L-N) E7 transgenic embryo showing native GFPcaax and nlsmCherry muscle specific expression of the transgene in all skeletal muscles.

This quail line is the first avian transgenic line dedicated to skeletal muscles and it should be useful to all researchers in the field interested in characterizing the dynamics of myogenic differentiation during embryogenesis.

#### CAAGS Kaede: ubiquitous expression of a photoactivatable fluorescent protein

Kaede is a photo-convertible protein (Ando et al., 2002) that undergoes irreversible photo-conversion from green to red fluorescence upon irradiation with violet or ultraviolet light. This fluorescent protein has been introduced in various animal models (mice, zebrafish) and is used in developmental biology as “optical cell marker” for short-term cell tracking. We generated a transgenic line, expressing Kaede, driven by an ubiquitous CAGGS promoter. A stable line was obtained in which strong and widespread expression of the photo-activatable fluorescent protein is observed in adult (Fig4 A) and developing embryos (Fig4 B). After 405nm UV laser illumination of a region of interest (dotted lines in Fig4 C), the green Kaede photo-converts to red Kaede (Fig4 D-F) to optically label the cells.

**Figure 4.**
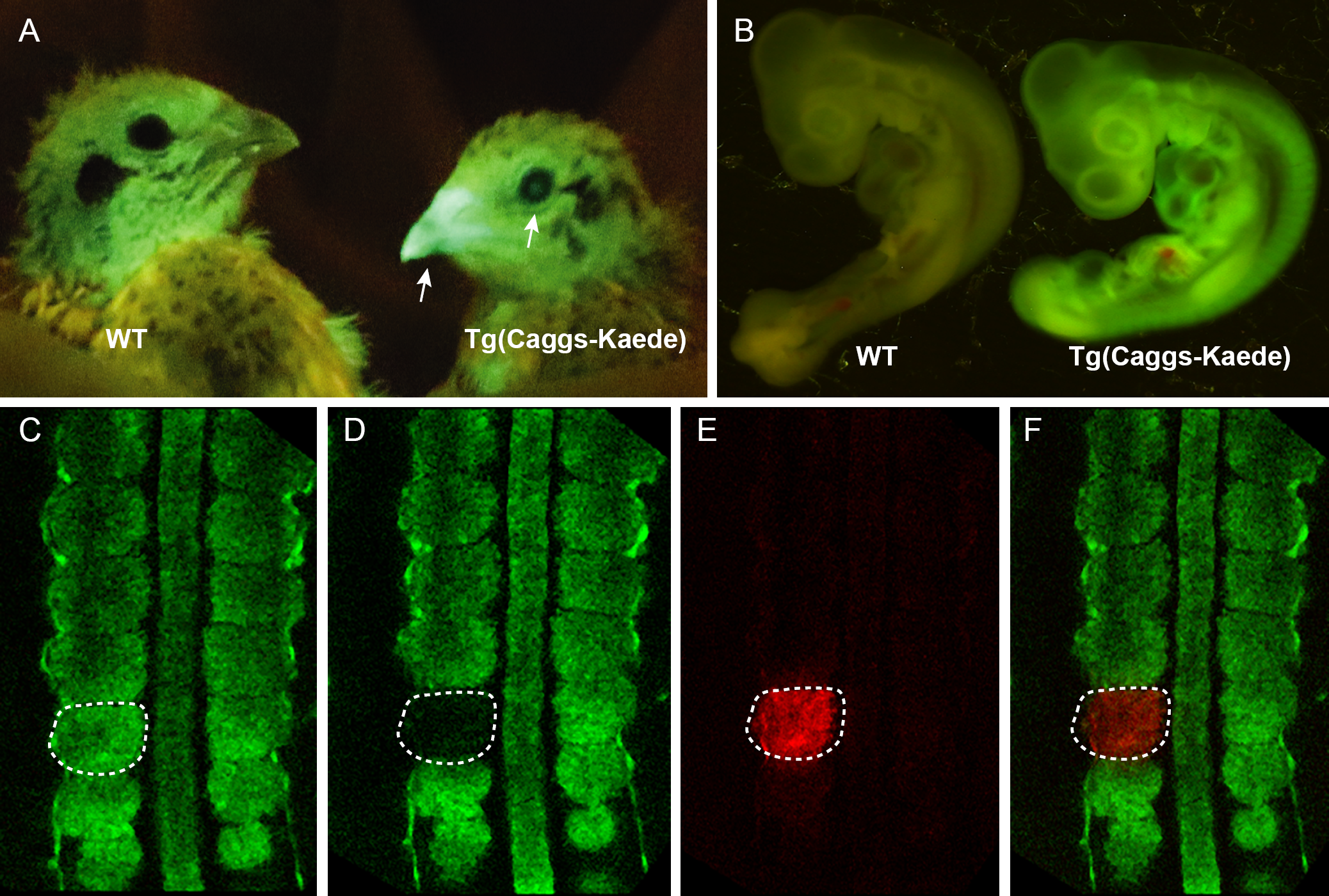
Description of the Tg(Caggs-Kaede) transgenic quail. Kaede expression is ubiquitously expressed in adult (A, arrows) and embryo (B). (C-D) using confocal 405nm UV laser and Region of Interest (ROI) tool, a specific area is illuminated (doted line) by UV laser (A) converting the Kaede protein present in the cells from green to red (D-F).

This quail line is the first avian line carrying a photo-convertible fluorescent protein and it represents a unique tool to perform short to medium-term lineage tracing of cells as development proceeds.

## Discussion

Avian embryos have been a model of choice in developmental biology due to its remarkable similarities with the human embryo and its accessibility to micro-surgery, tissue grafting experiments or, more recently, *in vivo* electroporation. The amenability of avian embryos to direct observation also allowed in vivo time-lapse imaging of cell behaviour as development proceeds. However, its use has been hampered by the difficulty to modify the bird genome using the genetic approaches developed in other models. Recent years have seen the emergence of transgenic chicken, generated by infection with lentiviruses or by injection of genetically manipulated PGCs. While these methods have proven efficient, they present serious limitations. The plasmid-based, direct transfection of PGCs described here combines the simplicity of DNA preparation with few limitations in construct sizes, a high efficiency. Moreover, the quail short generation time, their small size and the large amount of eggs they lay represent an economic alternative to other vertebrate models. Future development of the plasmid-based, direct transfection approach is the generation of knock-out mutants using CRISPR. This completes the panel of classical and cutting-edge technologies that can be applied to the avian model, making it one of the most versatile experimental systems available.

## Material and methods

### Generating transgenic quail by direct injection

The direct injection technique is performed as described in (Tyack et al., 2013). Injection mix contained 0.6ug of Tol2 plasmid, 1.2ug of CAGGS Transposase plasmid, 3ul of lipofectamin 2000 in 90 ul of Optipro. 1ul of injection mix in injected in dorsal aorta of 2.5 day old embryo. After injection, eggs are sealed and incubated until hatching. Hatchlings were grown for 6 weeks until they reach sexual maturity. Semen from males was collected using a female teaser and the massage technique as described in (Chelmonska et al., 2008). The genomic DNA from semen was extracted and PCR was performed to test for the presence of the transgene in semen. Males showing a positive band in semen DNA were crossed with wild type females. Offspring were selected directly after hatching by using UV illumination and goggles if the expression of the transgene was readily visible in hatchlings or by genotyping 5 days after hatching by plucking a feather.

### Section, immunochemistry and confocal analyses

Embryos were dissected under fluorescent stereomicroscope and fixed for 1h in 4% formaldehyde. Embryos were embedded in 15% sucrose/7.5% gelatine/PBS solution and sectioned with a cryostat at 20µm. Antibody stainings were performed as described in (Serralbo and Marcelle, 2014). The following antibodies were used: anti-GFP chicken polyclonal (ab13970, Abcam;1/1500), anti-RFP rabbit polyclonal (ab6234, Abcam; 1/500), anti-Pax7 IgG1 mouse monoclonal (Hybridoma Bank; 1/10), anti-Vasa Rabbit polyclonal. Stained sections were examined using Leica SP5 confocal microscope. 20X immersion lens was combined with tile scan acquisition.

## Acknowledgment

The authors acknowledge Terry wise, Chris Darcy and Mark Tizard from CSIRO AAHL (Geelong Australia) for their expertise in direct injection and transgenic chicken breeding. The authors acknowledge Monash Micro Imaging, Monash University, for the provision of instrumentation, training and technical support. The Australian Regenerative Medicine Institute is supported by grants from the State Government of Victoria and the Australian Government.

